# Using karyotyping and achene morphology to accurately identify *Sonchus oleraceus* (Asteraceae) in the context of its biological control

**DOI:** 10.1101/2020.02.26.965863

**Authors:** M. Ollivier, P. Gauthier, V. Lesieur, T. Thomann, M. Jourdan, M. C. Bon, A. Sheppard, S. Raghu, M. S. Tixier, J.-F. Martin

## Abstract

In weed biocontrol, accurate identification of the target weed is essential to select effective and host-specific biocontrol agents. This study focuses on the biocontrol of an invasive weed species in Australia, native from Europe, the common sowthistle, *Sonchus oleraceus* L. (Asteraceae). During field surveys, the distinction between *S. oleraceus* and a morphologically related species (*S. asper* L.) was difficult because of specimens bearing intermediate morphological features. These observations raised questions about the reliability the morphological characters used for distinguishing between these species and the identity of the intermediate phenotypes. First, cytological analyses coupled with morphological comparisons were carried out on specimens collected in Europe and Australia. Results showed that specimens morphologically described as *S. oleraceus* and *S. asper* possessed, in accordance with literature 32 and 18 chromosomes, respectively. Specimens with intermediate morphotypes had 32 chromosomes, showing that they belong to *S. oleraceus* species. The variability of characters used for diagnosis is discussed and for a particular feature, achene ornamentation, an inquiry among 30 people was carried out to determine how this character might be relevant for distinguishing the two species herein considered. The successful identification rate was 92.2% (SE ± 0.77) showing the practical interest of this feature for diagnosis.

## Introduction

*Sonchus oleraceus* L. (Asteraceae: Cichorieae; common sowthistle) is a pioneer plant species, that colonises disturbed ecosystems, as roadsides, gardens, crops and fallows (Hutchinson et al. 1984). It is an annual, sometimes biennial, species (Holm et al. 1977), tolerant of a large range of temperatures (Chauhan et al. 2006). *Sonchus oleraceus* is considered native to Europe (Gleason and Cronquist 1991), North Africa and West Asia (Peschken 1982), but has subsequently spread to North and South America, East Asia and Oceania (Holm et al. 1977; Mejías and Andrés 2004). Common sowthistle has probably been introduced inadvertently in the late 18th century to Australia (Boulos 1974). It is now one of the most difficult weeds to control across some 4.3 M ha of crops in southeast Queensland and northern New South Wales (Walker et al. 2005; Osten et al. 2007) where it causes annual losses estimated at AUD$6.3M (Llewellyn et al. 2016). This weed has been managed using herbicides but due to the rapid development of multiple resistances, herbicide efficiency is decreasing (Boutsalis and Powles 1995; Adkins et al. 1997; Cook et al. 2014; Meulen et al. 2016; Llewellyn et al. 2016). A program of classical biological control was initiated for the integrated management of common sowthistle (McCarren and Scott 2008, 2013, 2017). This program clearly relies on the correct identification of the target plant to search for specialist natural enemies to manage focal taxa (Goeden and Ricker 1985; O’Hanlon et al. 2000; McFadyen 2003; Smith et al. 2010, 2018).

In surveys carried out in France, accurate identification of *S. oleraceus* was always obvious, using the diagnostic morphological characters commonly defined in literature, i.e. cauline leaf, auricle shape and achene ornamentation (Boulos 1974; Hutchinson et al. 1984; Gleason and Cronquist 1991). Some plants with intermediate morphological features of *Sonchus oleraceus* L. and *Sonchus asper* L. were observed. Such intermediate phenotypes questioned i) potential hybridization events, ii) the use of inappropriate morphological features to distinguish between the two species. In comparable research programs (e.g. *Senecio madagascariensis* (Scott et al. 1998), *Rhododendron ponticum* (Milne and Abbott 2000), *Acacia nilotica* (Wardill et al. 2005), *Rubus niveus* (St. Quinton et al. 2011), and *Rubus* spp. complex (Bruckart et al. 2017)), the need for clarification of the target plant identity has been recognized as a prerequisite for subsequent steps. Indeed, the specificity relationship between the host plant and its natural enemies might be very tight, e.g. for pathogens and Eriophyidae spider mites, which is a crucial element in the science of weed biocontrol (McFadyen 2003; Smith et al. 2010, 2018). Misidentification of the target weed and selection of unadapted agents can cause incomplete control or failure of the establishment of the control agent in invasive range (Goeden and Ricker 1985; O’Hanlon et al. 2000).

The first objective of the present study was to determine the status of the intermediate phenotypes observed. Both *S. oleraceus* and *S. asper* are self-compatible but can occasionally cross-pollinate, producing rare hybrid specimens with an intermediate number of chromosomes (hybrid, 2n = 25 chromosomes; *S. oleraceus*, 2n = 32; *S. asper*, 2n = 18 chromosomes) (Barber 1941; Walter and Kuta 1971; Hsieh et al. 1972; Mejías and Andrés 2004). It is commonly accepted that *Sonchus oleraceus* is a combination of *S. asper* and *Sonchus tenerrimus*. As an amphidiploid species, the genome of *S. oleraceus* integrates the genomic DNA of both parental species. Given that, the development of diagnostic molecular markers to delineate these species would be very complex (Kim et al. 2007), we thus choose use chromosome counts to clarify the status of the morphologically ambiguous specimens observed (Jauzein and Nawrot 2013; Pico and Dematteis 2014).

The second objective was to determine reliable morphological characters for distinguishing between *S. oleraceus* and *S. asper*. An inquiry was carried out to determine the easy use of one “simple” feature (i.e. achene ornamentation and shape) found to be an unambiguous character.

## Materials and Methods

### Description based on classical morphological features

In 2017 and 2018, during surveys carried out in France we had some difficulties to clearly distinguish *S. oleraceus* and *S. asper* individuals in the field. We thus sampled a total of 19 individuals among ten locations in Europe, but also five locations in Australia (to determine if same difficulties exist there) (Table 1), in areas known to be favourable for Sonchus sp., i.e. disturbed ecosystem as roadsides, gardens, crops and fallows (Hutchinson et al. 1984). Each specimen was characterized morphologically using the characters cited as the most significant discriminant between the species considered (Table 2): cauline leaf (degree of incision, margins serration, and aspect), auricle (shape and prominence) and achene (surface ornamentation). Other characters were not included in the description as they were not sufficiently variable to differentiate the species considered in the study (e.g. rosette leaves and stem traits) or because their observation is only possible during a limited period (e.g. 4-leaf stage, flowers that are in bloom in the early morning hours). A qualitative description was chosen as it fits the need for quick identification in the field (no need to perform measurements). Seeds from each specimen were collected and stored in paper envelopes with silica gel at ambient temperature (22°C) until further study.

**Table 1.**
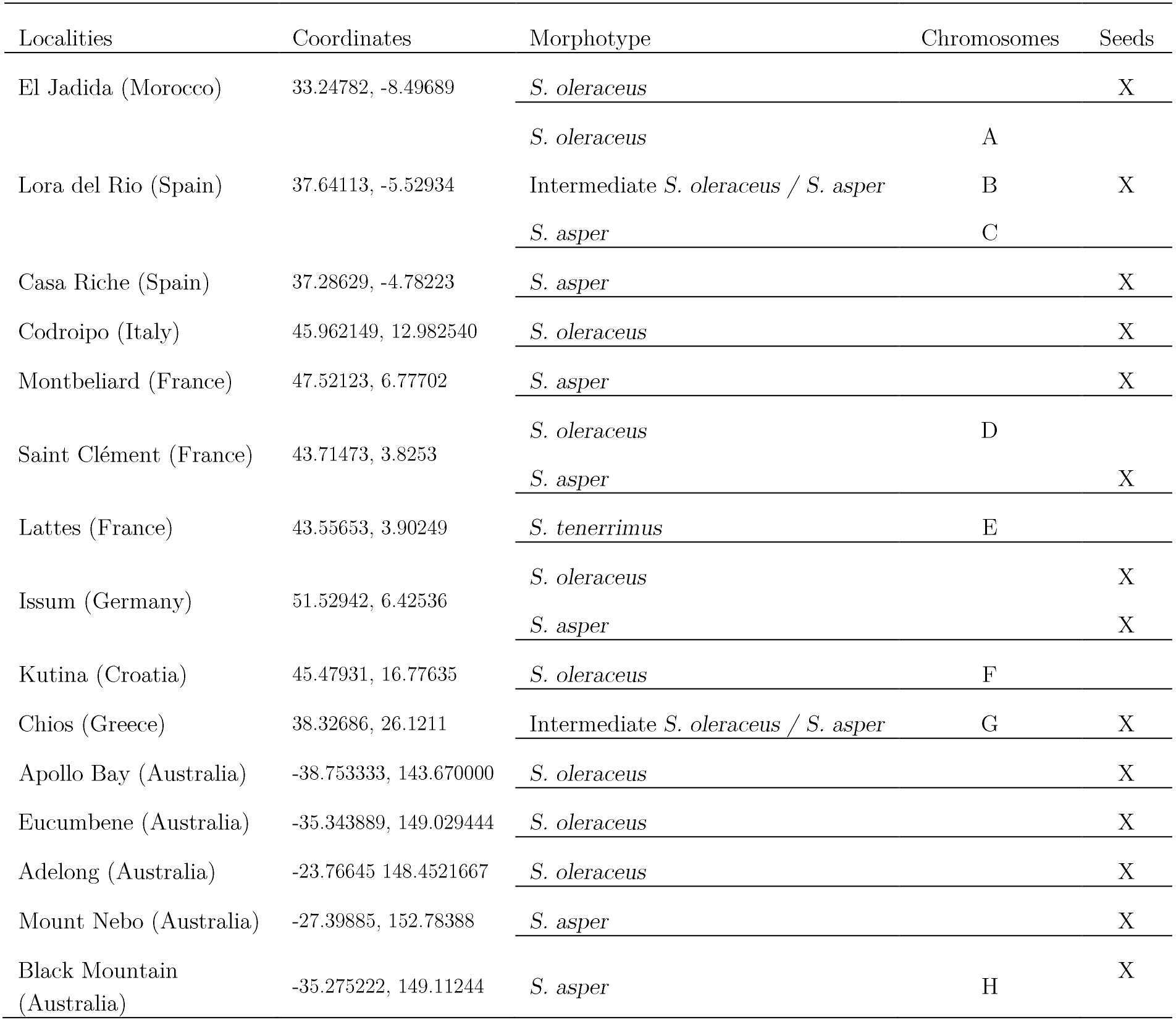
Localities where the different *Sonchus* species and morphotypes were collected in Europe and Australia, and selected for chromosome count and identification using achene ornamentation.

**Table 2:**
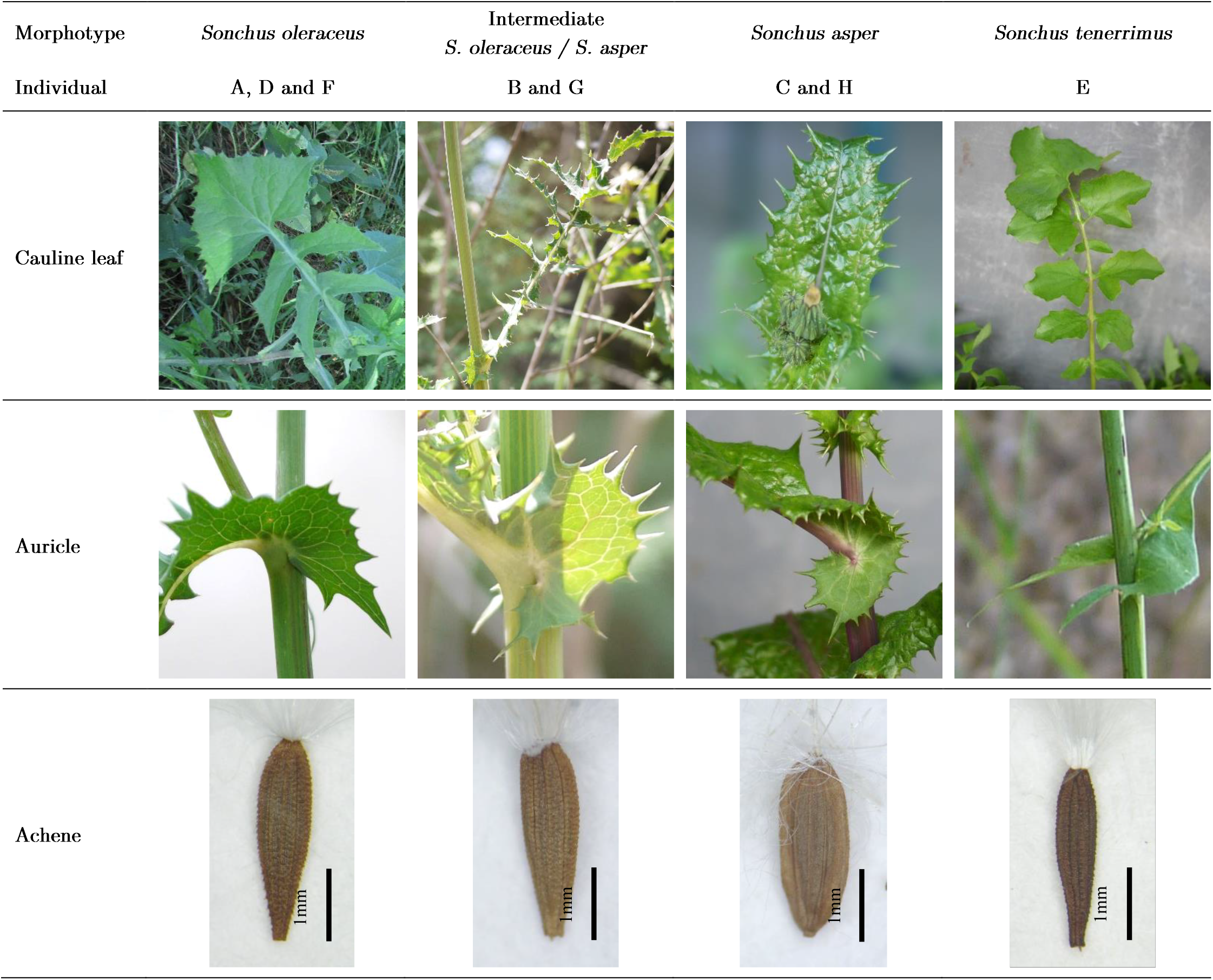
Marphological features (cauline leaves, auricles and surface ornamentation of achenes) for the speciemens collected.

| Morphotype | *Sonchus oleraceus* | Intermediate<br>*S. oleraceus* / *S. asper* | *Sonchus asper* | *Sonchus tenerrimus* |
| --- | --- | --- | --- | --- |
| Individual | A, D and F | B and G | C and H | E |
| Cauline leaf |  |  |  |  |
| Auricle |  |  |  |  |
| Achene |  |  |  |  |

### Mitotic Chromosome counting

#### Specimens considered

Among the 19 individuals, we chose eight specimens as representative of the four morphotypes of interest: (i) three individuals attributed to *S. oleraceus* (A, D, F), (ii) two individuals to *S. asper* (C, H), (iii) one individual to *S. tenerrimus* (E) and (iv) two ambiguous specimens (B, G) (table 1).

#### Karyotyping protocol

Mitotic chromosome counts were made from cells of young root tips that represent the most convenient source of mitotic cells (Doležel et al. 1999). A critical issue for chromosome counts is to obtain i) a sufficient number of well spread metaphase plates and a clear physical separation of the chromosomes and ii) a sufficient staining of the chromosomes owing to the small size of the chromosomes in Sonchus spp. This was achieved by optimizing the methods of Mirzaghaderi (2010). Cytological procedures have to be adapted as responses to pre-treatment, enzyme digestion conditions (concentration and incubation duration) are highly variable according to the species studied (Mirzaghaderi 2010; Kirov et al. 2014). Two key parameters were herein tested: (i) the period of the day at which the root tips are harvested and (ii) the duration of the immersion of the root tips into the enzymatic solution for cell wall digestion. Harvest time of root tips should correspond to the peak of the metaphase, that might vary over the day, as observed in other taxa as Euphorbia esula (Augé et al. 2016) and edible cocoyams (Ekanem and Osuji 2006). The harvest times of root tips tested were 8h00, 10h00 and 12h00. Immersion times tested were 45, 50, 55, 60, 65 and 70 minutes. The impact of each factor on the number of cells in division per meristem and the number of cells for which chromosomes are sufficiently spread to be counted over the total cells in division was tested separately with a one-way parametric ANOVA followed by Tukey pairwise comparisons using R3.4.1 (R Core Team 2017). The period of the day was tested on specimens A and D, with two repetitions per treatment. The immersion duration in enzymatic solution was tested on specimen D, with two repetitions per treatment.

#### Final protocol used

Germinated seedlings were obtained from seeds collected in the field. Seeds were set on wet Whatman ^®^ filter paper covering a substrate of soaked vermiculite. Seeds were maintained at temperature 25°C/20°C (day/night) to trigger germination. Three days-old root tips of 2-4 mm long were harvested at 12h00 (see results of protocol adjustment) and treated with 0.04% 8-hydroxyquinoline for 4 hours at room temperature, fixed in Farmer’s fixative solution ethanol: acetic acid (3:1) for 48 hours at room temperature, and stored at 4°C in 70% ethanol. The following day successive immersion in 0.25M HCL, citric buffer (pH 4.5) and enzyme digestion mixture (Cellulase Onozuka, Pectolyase Y-23 and Cytohelicase, 1:1:1) for one hour (see results of protocol adjustment) at 37°C in incubator were necessary to digest cell wall. Digested tissues were maintained during 2 hours in distilled water at ambient temperature, and then were gently transferred to a slide. A drop of fresh and cold (4°C) Farmer’s fixative solution ethanol:acetic acid (3:1) was added to improve chromosomes spread. Once the slide dry, DAPI (4’-6-diamino-2phenylinole) fluorochrome solution was applied and slide was left fifteen minutes in dark environment for DAPI to bind with DNA. VECTASHIELD ^®^ Antifade Mounting Medium was employed to preserve fluorescence before covering the slide and strongly squashing to enhance spreading. The slides were examined using a fluorescence microscope (Zeiss Axioplan imager A1) at 100x magnification and the images were obtained with a Jai camera (Cv-m4 CL).

### Diagnostic character selection and assessment

According to chromosome count results, we reconsidered the morphological characters described in the literature to target the optimal feature for field diagnosis. Achene ornamentation seemed to be a straightforward and quick feature to identify the considered species in field conditions. To better assess its reliability, achenes from six clearly identifiable plants of *S. oleraceus* and *S. asper* were selected, as well as the achenes of the two plants with intermediates morphotypes (and previously identified as *S. oleraceus* using chromosome counts). Ten achenes of these 14 specimens (140 in total) were randomly chosen and set on an A4 sticker paper and scanned (Epson Perfection V550 Photo, 1200dpi). Thirty persons including 15 taxonomists and 15 less-experienced persons were asked to identify the 140 achenes, using the illustrative pictures presented in Table 2. The error rate in distinguishing the two species was calculated and standard error was estimated. The difference in scores of experienced and less-experiences participants was tested with a one-way parametric ANOVA. To investigate whether the misidentifications obtained were particularly attributed to achenes belonging to ambiguous specimens, a non-parametric Wilcoxon signed-rank test was carried out to compare score from ambiguous vs clearly identifiable specimens of *S. oleraceus* and *S. asper*. All analyses were run under R3.4.1 (R Core Team 2017).

## Results and discussion

### Description based on classical morphological features

Morphological characters displayed by the specimens collected are illustrated in Table 2. Nine specimens (among which are the specimens used in chromosome counts: A, D and F) presented the following features: cauline leaf shape was pinnatifid to whole limb, slightly to deeply lobed, with lobe base section superior to 5 mm long and leaf margin toothed but smoothed. The aspect of the cauline leaves was mat. Auricles were sharply acute and projected forward. Achene ornamentation showed two to five prominent ribs with transverse ridges. These features correspond to those of *S. oleraceus* (Boulos 1974; Hutchinson et al. 1984; Gleason and Cronquist 1991). Seven specimens (among which are the specimens used in chromosome countings: C and H) displayed cauline leaf fully to slightly lobed, with leaf margin toothed, spiny and a glossy aspect. Auricles were prominent, rounded, spiny and clasped the stem. Achenes displayed three prominent ribs but no transverse ridges. These morphological features match with those of *S. asper* (Hsieh et al. 1972; Boulos 1974). Two specimens (B and G) displayed “ambiguous” morphological characteristics, i.e. intermediate between *S. oleraceus* and *S. asper* (Table 2). Cauline leaves were pinnatifid to whole limb, slightly to deeply lobed, with weakly to hardly spinous leaf margins, and a mat to glossy aspect. Auricles were big, rounded, with projected teeth but not sharply acute, which could be associated to *S. asper*. In contrast, achenes showed two to five prominent ribs with transverse ridges, feature usually distinctive of *S. oleraceus*. Finally, the features of the specimen E correspond to those of the description of *S. tenerrimus* (Boulos 1974). Cauline leaves were pinnatifid, with various lobe shapes but the lobe base section was always inferior to 5 mm long. Auricles were sharply acute and projected forward, while achenes presented one to three prominent ribs with transverse ridges.

### Protocol development for counting chromosomes

For the Sonchus species considered, the optimal period of the day to harvest root tips was 12h00 as the number of cells in division per meristem and the proportion of cells with countable chromosomes were significantly higher than at 8h00 and 10h00 (mean of 2.75 cells in division at 8h00, 2.25 at 10h00 and 10 at 12h00, and mean proportion of 12.5% of countable cells at 8h00, 16.2% at 10h00 and 63.9% at 12h00; F(2, 9) = 5.3, p-value = 0.0311 and F(2,9) = 7.67, p-value = 0.0113 respectively). Growing conditions, such as light and temperature, are known to affect cell cycle (Doležel et al. 1999). Cell division is mediated by the circadian clock and usually occurs at specific times of the day in animal and plant organisms (Moulager et al. 2007). These results suggest that the best time to collect root tips for Sonchus sp. for optimum metaphase is around noon. Similar observations were made for different plant taxa grown under different conditions (Ekanem and Osuji 2006; Augé et al. 2016). The duration of enzymatic digestion did not have any significant effect on the quality of the observation (mean of 4.5 cells in division after 45min, 0 after 50min, 1 after 55 min, 0.5 after 60min, 0 after 65min and 4.5 after 70min; F(1, 10) = 0.001, p-value = 0.972), hence one hour was the duration used for all the cytological experiments.

### Chromosome counts

Results of chromosome counts are presented in Table 3. The *S. oleraceus* specimens (A, D and F) had 2n=32 chromosomes, the *S. asper* specimens (C and H) displayed 2n=18 chromosomes and the *S. tenerrimus* specimen (E) had 2n=14 chromosomes. These results are consistent with literature (Walter & Kuta 1971; Hsieh et al. 1972; Boulos 1974; Mejías & Andrés 2004). For the specimens with intermediate morphotypes (B and G), the chromosome count was 2n=32 chromosomes, suggesting that these specimens belong to *S. oleraceus*. They are not hybrids between *S. oleraceus* and *S. asper* for which Barber (1941) counted 2n=25 chromosomes.

**Table 3:**
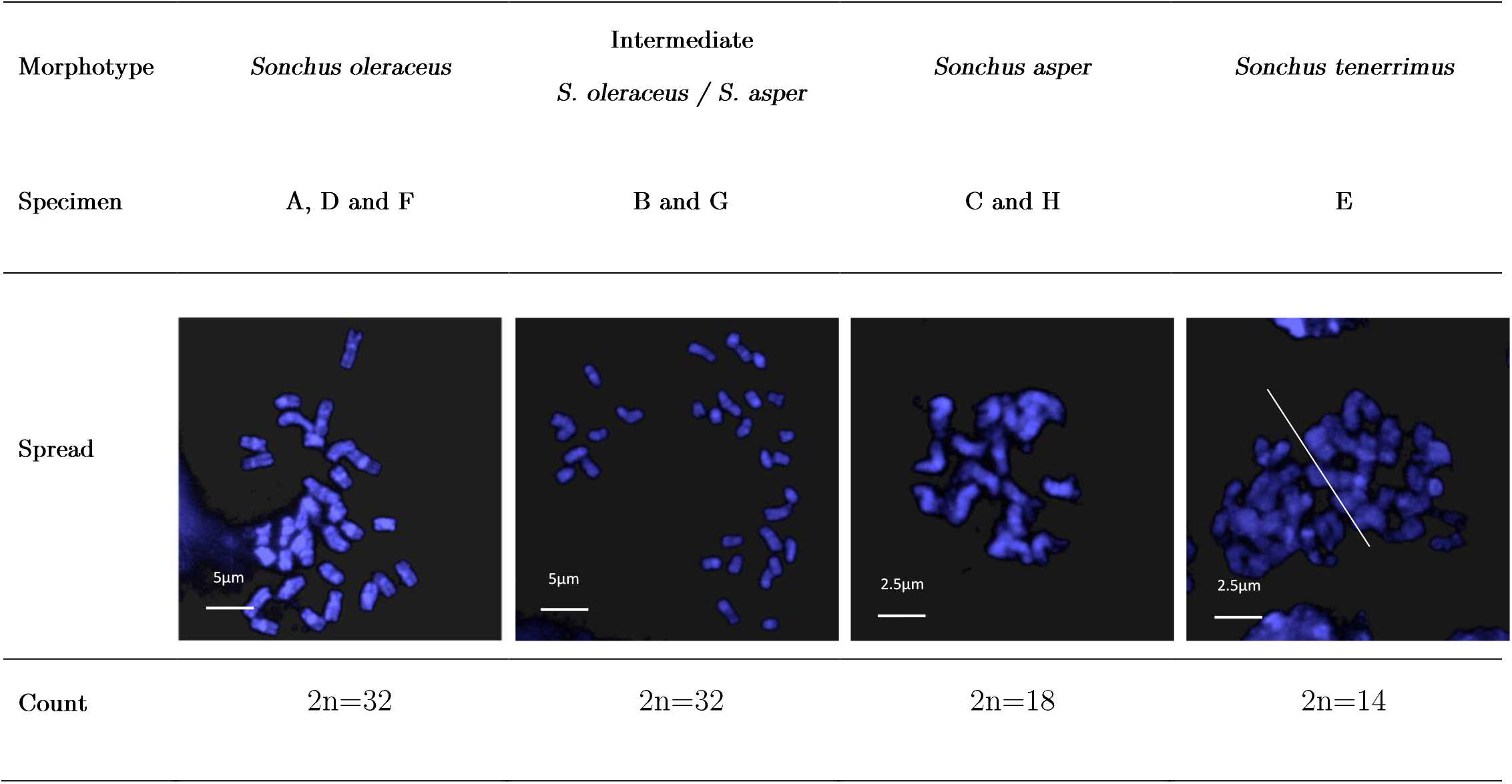
Mitotic chromosome spread and count obtained at metaphase stage from root tip meristems of the four different *Sonchus* morphotypes. (DAPI staining, at x100 magnification)

| Morphotype | *Sonchus oleraceus* | Intermediate<br>*S. oleraceus* / *S. asper* | *Sonchus asper* | *Sonchus tenerrimus* |
| --- | --- | --- | --- | --- |
| Specimen | A, D and F | B and G | C and H | E |
| Spread |  |  |  |  |
| Count | 2n=32 | 2n=32 | 2n=18 | 2n=14 |

The ambiguous morphotypes underline the high variability in morphological traits of *S. oleraceus* (Boulos 1974; Gleason and Cronquist 1991; Widderick 2002; Widderick et al. 2004; Jauzein and Nawrot 2013). This result questions the reliable use of cauline leaves and auricles shape for distinguishing between *S. oleraceus* and *S. asper*.

### Defining a reliable diagnostic character for field identification

Although cauline leaves are easy to observe at any time (except the rosette stage), ornamental features of achenes seem more reliable. The achene ornamentation of *S. oleraceus* includes two to five prominent ribs with transverse ridges (Table 2) (Boulos 1974; Hutchinson et al. 1984; Gleason and Cronquist 1991). The presence of transverse wrinkles for *S. oleraceus* (specimens A, B, D, F and G) clearly contrasts with the three prominent and smooth ribs displayed on *S. asper* achenes (specimens C and H) and represent a clear diagnostic feature. In addition, Boulos (1973) reported flowers as reliable traits to separate *S. oleraceus*, *S. asper* and *S. tenerrimus*. He proposed the ratio between ligule and corolla tube lengths whose value for the species cited are respectively 1:1, 2:3 and 4:3. However, flowers only open early in the morning and are fully observable on the plant only at late phenological stages, which might hinder the number of surveys required for searching for natural enemies on an unambiguously correct targeted weed species. In contrast to flower and seedling features, achene ornamentation is accessible for a longer period. Before flower bloom and fruit complete formation, striations are already observable and once the flower opened and fruits fully developed a proportion of achenes remain in the flower head for several weeks (one to four weeks, depending on abiotic conditions), entangled among pappus.

Distinction between *S. asper* and *S. oleraceus* seems at first sight straightforward and rapid, simply using a field magnifier. To determine if this latter hypothesis is right, we tested the reliability of this trait on a subset of the plants used in previous stages (see material and methods) on a group of 30 participants (half of them were used to species delineation (entomologist) and half had low to no experience). The successful identification rate of species based on achene ornamentation was 92.2% (SE ± 0.77). A significant difference was observed between the two people groups (F(1, 28) = 10.56, p-value = 0.003) due to their contrasted habits in species identification. Although experienced taxonomist reached a higher success rate (94.4%, SE ± 0.70) than novice participants (90.0%, SE ± 1.13), this score is readily acceptable considering the lack of botanic skills of the people evaluated. Feature validation are rarely realised (Balakrishnan 2005), so these scores are difficult to compare with similar studies, but both were deemed highly satisfying. Contrary to Austen et al. (2016) who gave the persons the possibility to not respond in case of doubts, in our survey, people were forced to provide an answer. Likewise, using an image instead of fresh material observation can lead to mistakes, as people could not angle the achene and adjust the focus as they wish. The errors herein observed were not particularly attributed to achenes from ambiguous specimens, as they were in average less misidentified than other specimens (94.8% SE ± 1.42 and 91.8% ± SE 0.76, of success respectively / W = 238.5, p-value = 0.001). A discussion with some of the participants allowed to get feedback on the identification: beyond the contrast in rugosity of the achene face, the global shape of the achene was the most helpful criterion for discrimination. *S. oleraceus* elongated shape is promptly distinguished from *S. asper* oblong shape. In case of any doubt for some achenes, it should be notice that in field conditions additional achenes belonging the same flower displaying more pronounced traits could be sampled. Here, we therefore confirm the relevance of achene ornamentation as a diagnostic character.

## Conclusion

Chromosome counting allowed refuting the hybridization hypothesis to explain the occurrence of the morphological intermediate specimens collected and confirmed the high degree of morphological variation within *S. oleraceus* probably enhanced by its amphidiploid origin. The low reliability of several plant features currently used to identify *S. oleraceus* and *S. asper* (e.g. cauline leaves and auricles) may cause misidentification of the target weed in Europe with consequences for the selection of biocontrol agents (O’Hanlon et al. 2000). The high success rate in distinguishing *S. oleraceus* from *S. asper* (mean 92.2%, SE ± 0.77) obtained with a large group of expert to non-expert botanists, strongly supports the usefulness of achene ornamentation for the identification of these two Sonchus species in the field conditions. More generally, we can notice that identification of plants at late stage based on achene features is of interest for weed biocontrol as it enable to look for promising biocontrol agent belonging to seed feeder arthropods. Finally, clarifying Sonchus complex and determining relevant morphological features for their differentiation is essential for subsequent assessments of biocontrol agent establishment in Australia, the targeted area, where both *S. oleraceus* and *S. asper* coexist.

## Acknowledgments

Chromosome counting presented in the present publication were performed at the “Génomique Environnemental/Cytogénomique Evolutive” facilities from the LabEx CeMEB (Laboratoire d’Excellence Centre Méditerranéen de l’Environnement et de la Biodiversité, Montpellier, France). This project is supported by funding from the Australian Government Department of Agriculture and Water Resources, as part of its Rural R&D for Profit programme, through AgriFutures Australia (Rural Industries Research and Development Corporation) (PRJ--010527). Fatiha Guermache is gratefully acknowledged for technical assistance and her wise advice. Warm thanks are also addressed to all the participants of the identification test.

## References

Adkins, S.W., Wills, D., Boersma, M., Walker, S.R., Robinson, G., Mcleod, R.J., and Einam, J.P. 1997. Weeds resistant to chlorsulfuron and atrazine from the north-east grain region of Australia. Weed Res. 37(5): 343–349. doi:10.1046/j.1365-3180.1997.d01-56.x.

Augé, M., Bon, M.-C., Hardion, L., Le Bourgeois, T., and Sforza, R.F.H. 2016. Genetic characterization of a red colour morph of Euphorbia esula subsp. esula (Euphorbiaceae) in the floodplains of the Saône (Eastern France). Botany 94(10): 1001–1007. doi:10.1139/cjb-2016-0067.

Austen, G.E., Bindemann, M., Griffiths, R.A., and Roberts, D.L. 2016. Species identification by experts and non-experts: comparing images from field guides. Sci. Rep. 6: 33634. doi:10.1038/srep33634.

Balakrishnan, R. 2005. Species Concepts, Species Boundaries and Species Identification: A View from the Tropics. Syst. Biol. 54(4): 689–693. doi:10.1080/10635150590950308.

Boulos, L. 1974. Revision systematique du genre Sonchus L. s.l. VI. Sous-genre. 3. origosonchus. genres Embergeria, Babcockia et - Taeckholmia. species exclusae et dubiae. index. Bot. Not. Available from http://agris.fao.org/agris-search/search.do?recordID=US201303094110 [accessed 13 July 2017].

Boutsalis, P., and Powles, S. 1995. Resistance of dicot weeds to acetolactate synthase (ALS)-inhibiting herbicides in Australia. Weed Res. 35: 149–155. doi:10.1111/j.1365-3180.1995.tb02028.x.

Bruckart, W.L., Michael, J.L., Sochor, M., and Trávníček, B. 2017. Invasive Blackberry Species in Oregon: Their Identity and Susceptibility to Rust Disease and the Implications for Biological Control. Invasive Plant Sci. Manag. 10(2): 143–154. doi:10.1017/inp.2017.12.

Chauhan, B.S., Gill, G., and Preston, C. 2006. Factors affecting seed germination of annual sowthistle (*Sonchus oleraceus*) in southern Australia. Weed Sci. 54(5): 854–860. doi:10.1614/WS-06-047R.1.

Cook, T., Davidson, B., and Miller, R. 2014. A new glyphosate resistant weed species confirmed for northern New South Wales and the world: common sowthistle (*Sonchus oleraceus*). 19th Australas. Weeds Conf. Sci. Community Food Secur. Weed Chall. Hobart Tasman. Aust. 1-4 Sept. 2014: 206–209.

Doležel, J., Číhalíková, J., Weiserová, J., and Lucretti, S. 1999. Cell cycle synchronization in plant root meristems. Methods Cell Sci. 21(2): 95–107. doi:10.1023/A:1009876621187.

Ekanem, A.M., and Osuji, J.O. 2006. Mitotic index studies on edible cocoyams (Xanthosoma and Colocasia spp.). Afr. J. Biotechnol. 5(10). Available from https://www.ajol.info/index.php/ajb/article/view/42881 [accessed 17 September 2018].

Gleason, H.A., and Cronquist, A. 1991. Manual of Vascular Plants of Northeastern United States and Adjacent Canada. In Second. New York Botanical Garden Press. doi:10.21135/893273651.001.

Goeden, R.D., and Ricker, D.W. 1985. Seasonal Asynchrony of Italian Thistle, Carduus pycnocephalus, and the Weevil, Rhinocyllus conicus (Coleoptera: Curculionidae), Introduced for Biological Control in Southern California. Environ. Entomol. 14(4): 433–436. doi:10.1093/ee/14.4.433.

Holm, L.G., Plucknett, D.L., Pancho, J.V., and Herberger, J.P. 1977. The world’s worst weeds. Distribution and biology. Worlds Worst Weeds Distrib. Biol. Available from https://www.cabdirect.org/cabdirect/abstract/19782319336 [accessed 4 September 2018].

Hsieh, T.-S., Schooler, A.B., Bell, A., and Nalewaja, J.D. 1972. Cytotaxonomy of Three Sonchus Species. Am. J. Bot. 59(8): 789–796. doi:10.2307/2441083.

Hutchinson, I., Colosi, J., and Lewin, R.A. 1984. THE BIOLOGY OF CANADIAN WEEDS.: 63. *Sonchus asper* (L.) Hill and S. oleraceus L. Can. J. Plant Sci. 64(3): 731–744. doi:10.4141/cjps84-100.

Jauzein, P., and Nawrot, O. 2013. Flore d’Île-de-France: Clés de détermination, taxonomie, statuts. Editions Quae.

Kim, S.-C., Chunghee, L., and Mejías, J.A. 2007. Phylogenetic analysis of chloroplast DNA matK gene and ITS of nrDNA sequences reveals polyphyly of the genus Sonchus and new relationships among the subtribe Sonchinae (Asteraceae: Cichorieae). Mol. Phylogenet. Evol. 44(2): 578–597. doi:10.1016/j.ympev.2007.03.014.

Kirov, I., Divashuk, M., Van Laere, K., Soloviev, A., and Khrustaleva, L. 2014. An easy “SteamDrop” method for high quality plant chromosome preparation. Mol. Cytogenet. 7(1): 21. doi:10.1186/1755-8166-7-21.

McCarren, K.L., and Scott, J.K. 2008, March 24. Two biological control options for *Sonchus oleraceus* in Australia. Available from https://eurekamag.com/research/034/095/034095536.php [accessed 16 August 2017].

McCarren, K.L., and Scott, J.K. 2013. Host range and potential distribution of Aceria thalgi (Acari: Eriophyidae): a biological control agent for Sonchus species. Aust. J. Entomol. 52(4): 393–402. doi:10.1111/aen.12041.

McCarren, K.L., and Scott, J.K. 2017. Host range and potential distribution of the rust fungus, Miyagia pseudosphaeria, a biological control agent for Sonchus species. Australas. Plant Pathol. 46(5): 473–482. doi:10.1007/s13313-017-0509-9.

McFadyen, R.E. 2003. Does ecology help in the selection of biocontrol agents? Tech. Ser. - CRC Aust. Weed Manag. (No.7): 5–9.

Mejías, J.A., and Andrés, C. 2004. Karyological studies in IberianSonchus (Asteraceae: Lactuceae): *S. oleraceus*, S. microcephalus and *S. asper* and a general discussion. Folia Geobot. 39(3): 275–291. doi:10.1007/BF02804782.

Meulen, A.W. van der, Widderick, M., Cook, T., Chauhan, B.S., and Bell, K. 2016. Survey of glyphosate resistance in common sowthistle (*Sonchus oleraceus*) across the Australian Northern Grains Region. In 20th Australasian Weeds Conference. Perth, Western Australia. p. 108. Available from http://caws.org.au/awc/2016/awc201611081.pdf [accessed 30 March 2018].

Milne, R.I., and Abbott, R.J. 2000. Origin and evolution of invasive naturalized material of Rhododendron ponticum L. in the British isles. Mol. Ecol. 9(5): 541–556.

Mirzaghaderi, G. 2010. simple metaphase chromosome preparation from meristematic root tip cells of wheat for karyotyping or in situ hybridization. Afr. J. Biotechnol. 9(3). Available from https://www.ajol.info/index.php/ajb/article/view/77903 [accessed 5 April 2018].

Moulager, M., Monnier, A., Jesson, B., Bouvet, R., Mosser, J., Schwartz, C., Garnier, L., Corellou, F., and Bouget, F.-Y. 2007. Light-Dependent Regulation of Cell Division in Ostreococcus: Evidence for a Major Transcriptional Input. Plant Physiol. 144(3): 1360–1369. doi:10.1104/pp.107.096149.

O’Hanlon, P., Briese, D., and Peakall, R. 2000. Know Your Enemy: The Use of Molecular Ecology in the Onopordum Biological Control Project. In Proceedings of the X International Symposium on Biological Control of Weeds. Advanced Litho Printing. Available from https://openresearch-repository.anu.edu.au/handle/1885/91055 [accessed 23 May 2017].

Osten, V.A., Walker, S.R., Storrie, A., Widderick, M., Moylan, P., Robinson, G.R., and Galea, K. 2007. Survey of weed flora and management relative to cropping practices in the north-eastern grain region of Australia. Aust. J. Exp. Agric. 47(1): 57–70. doi:10.1071/EA05141.

Peschken, D.P. 1982. Host specificity and biology ofcystiphora sonchi [Dip.: Cecidomyiidae], a candidate for the biological control ofSonchus species. Entomophaga 27(4): 405–415. doi:10.1007/BF02372063.

Pico, G.M.V.D., and Dematteis, M. 2014. Cytotaxonomy of two species of genus Chrysolaena H. Robinson, 1988 (Vernonieae, Asteraceae) from Northeast Paraguay. Comp. Cytogenet. 8(2): 125–137. doi:10.3897/CompCytogen.v8i2.7209.

Scott, L.J., Congdon, B.C., and Playford, J. 1998. Molecular evidence that fireweed (Senecio madagascariensis, Asteraceae) is of South African origin. Plant Syst. Evol. 213(3-4): 251–257. doi:10.1007/BF00985204.

Smith, L., Cristofaro, M., Bon, M.-C., De Biase, A., Petanović, R., and Vidović, B. 2018. The importance of cryptic species and subspecific populations in classic biological control of weeds: a North American perspective. BioControl 63(3): 417–425. doi:10.1007/s10526-017-9859-z.

Smith, L., de Lillo, E., and Amrine, J.W. 2010. Effectiveness of eriophyid mites for biological control of weedy plants and challenges for future research. Exp. Appl. Acarol. 51(1-3): 115–149. doi:10.1007/s10493-009-9299-2.

St. Quinton, J.M., Fay, M.F., Ingrouille, M., and Faull, J. 2011. Characterisation of Rubus niveus⍰z: a prerequisite to its biological control in oceanic islands. Biocontrol Sci. Technol. 21(6): 733–752. doi:10.1080/09583157.2011.570429.

Walker, S.R., Taylor, I.N., Milne, G., Osten, V.A., Hoque, Z., and Farquharson, R.J. 2005. A survey of management and economic impact of weeds in dryland cotton cropping systems of subtropical Australia. Aust. J. Exp. Agric. 45(1): 79–91. doi:10.1071/EA03189.

Walter, R., and Kuta, E. 1971. Cytological and embryological studies in Sonchus L. I. *Sonchus asper* (L.) Hill. and *Sonchus oleraceus* L. Acta Biol. Cracoviensia Ser. Bot. Available from http://agris.fao.org/agris-search/search.do?recordID=US201302246800 [accessed 4 April 2018].

Wardill, T.J., Graham, G.C., Zalucki, M., Palmer, W.A., Playford, J., and Scott, K.D. 2005. The importance of species identity in the biocontrol process: identifying the subspecies of Acacia nilotica (Leguminosae: Mimosoideae) by genetic distance and the implications for biological control. J. Biogeogr. 32(12): 2145–2159. doi:10.1111/j.1365-2699.2005.01348.x.

Widderick, M., Walker, S., and Sindel, B. 2004. Better management of *Sonchus oleraceus* L. (common sowthistle) based on the weed’s ecology. Weed Manag. Balanc. People Planet Profit 14th Aust. Weeds Conf. Wagga Wagga New South Wales Aust. 6-9 Sept. 2004 Pap. Proc.: 535–537.

Widderick, M.J. 2002. Ecology and Management of the Weed Common Sowthistle (*Sonchus oleraceus* L.). University of New England.

